# Genomic analysis in *Entamoeba* reveals intron gain and unbiased intron loss, transformed splicing signals, and a coevolved snRNA

**DOI:** 10.1101/2022.06.08.495308

**Authors:** Scott William Roy, Bradley A. Bowser

## Abstract

The intron-exon structures of nuclear genes show striking diversity across eukaryotes. Several independent lineages have undergone convergent evolution including widespread loss of introns and transformed *cis* splicing signals. The causes and mechanisms of these changes remain mysterious: (i) transformation of splicing signals could reflect either selective loss of suboptimal introns or coevolution of introns and splicing machinery; and (ii) corresponding changes in the splicing machinery remain poorly characterized. A promising model to study these questions is *Entamoeba.* Analysis of five *Entamoeba* species revealed low intron densities, nearly universal atypical 5’ splice sites and 3’ intronic sequences. A flexible search for U1 snRNA genes revealed a modified 5’-AACAAAC-3’ recognition sequence, affording complete Watson-Crick basepairing potential with the atypical 5’ splice site and extended basepairing potential. A U1 candidate in the related species *Mastigamoeba balumuthi* revealed a separate modification complimenting a different atypical consensus splice site. Genome-wide study of intron loss and gain revealed that introns with suboptimal splicing motifs were no more likely to be lost, suggesting against genome-wide homogenization of intron splicing motifs by selective intron loss. Unexpectedly, this analysis also revealed widespread intron gain in *Entamoeba invadens*. In total, the current analyses: (i) provide the most direct available evidence of coevolution of spliceosomal introns and splicing machinery; (ii) illuminate the evolutionary forces responsible for concerted intron loss and splicing motif transformation; and (iii) reveal widespread intron gain in an otherwise highly reduced lineage.

## Introduction

In most characterized eukaryotic organisms, the majority of protein-coding transcripts are processed by a large RNA-protein machinery termed the spliceosome, which removes internal portions of transcripts, termed spliceosomal introns, which are thus not represented in mature mRNA transcripts. The genomic era has revealed the ubiquity of spliceosomal introns within eukaryotes, with the genomes of nearly all characterized eukaryotes encoding spliceosomal components and containing spliceosomal introns (1–5).

At the same time, comparative genomic studies have revealed striking differences in the characteristics of spliceosomal systems across eukaryotes, indicating a dynamic evolutionary history. Striking differences across eukaryotes are seen in the numbers, splicing signals and lengths of spliceosomal introns, as well as in in the spliceosomal itself (3). Intron number range in numbers from hundreds of thousands in vertebrates to apparently only one in some kinetoplastids (6–7); in lengths from a broad distribution with a mean of five kilobases in humans to nearly all introns being 15 or 16 nts in *Stentor coeruleus* (6,8); in splicing motifs from the diverse motifs dispersed across exons and introns characteristic of many species to highly homogeneous concentrated core intron sequences in species such as yeast (9–11); and in complexity of machinery from hundreds of spliceosomal proteins in animals to perhaps a few dozen in the red alga *Cyanidioschyzon merolae*, which appears to have entirely lost the U1 snRNP, the one of the five core complexes of the spliceosome responsible for recognizing the 5’ splice site (12–15).

A variety of studies have provided the outlines of this dynamic evolutionary history. Evolutionary reconstructions have revealed that early eukaryotic ancestors harbored a complex spliceosomal system, with large numbers of introns, complex dispersed splicing signals, and a complex spliceosome (9,12,16–23). Thus, lineages exhibiting fewer introns, simpler splicing signals, or simplified spliceosomal machinery have undergone simplification. Comparative analysis of auxiliary exonic signals has suggested that such ‘simplified’ species also differ from other species in greatly reduced usage of such signals, suggesting recurrent evolution from diffuse splicing signals relying on diverse components to more concentrated signals relying largely on core splicing motifs (10,24). Notably, such simplification has occurred many times in eukaryotic history. For instance, lineages from within animals, unicellular holozoans, microsporidia, apicomplexans, red algae, green algae, diplomonads, parabasalids, apusozoa, animals and amoebozoa have independently undergone widespread reduction in intron number, leading to clear minorities of genes in these reduced lineages harboring introns, in each case representing at least an order-of-magnitude reduction relative to ancestral genomes (3,9,23). This extensive sampling of eukaryotic genomes also allows for tracing the coevolution of different traits of the spliceosomal system. In some cases, correspondences have been found that fit many scientists’ intuitive expectation: for instance, simplification of the splicing machinery seems to have occurred in lineages with very few introns (14,25–26). In other cases, clear but unexpected correspondences have been discovered: for instance, all lineages that have undergone widespread reduction in intron number have also undergone homogenization of splicing signals, with introns in these genomes showing highly homogeneous splicing signals (for instance, three-quarters of introns in the yeast *S. cerevisiae* have the 5’ splice site GUAUGU (3,9,23)). In still other cases, correspondences that might be expected intuitively have not been recovered. In particular, although changes in splicing signals imply differences in intron recognition mechanisms (since if the observed signals were not required for efficient recognition of introns, we would expect ongoing mutation to produce a diversity of splice sites), corresponding changes in the splicing machinery have been elusive.

One particular case of transformation remains mysterious. In most modern species as well as reconstructed ancestors, 5’ splice sites of introns show a consensus sequence of 5’-GUAAGU-3’, which provides complete complementarity to the corresponding region of the U1 snRNA (5’-ACUUAC-3’), which is consistent with the U1 snRNA’s role in splice site recognition through basepairing with the splice site. Interestingly, a variety of lineages, which appear to be concentrated among those that have undergone reduction in intron number, have evolved atypical 5’ splice sites, with a clear consensus of 5’-GUAUGU-3’ ((3) and unpublished data). However, these lineages have not evolved a complementary change in their U1 snRNA, as might be expected given the importance of basepairing for intron recognition (27). This lack of complementary change has been found in multiple lineages favoring 5’-GUAUGU-3’ splice sites, including Saccharomycotina yeasts, diplomonads such as *Giardia lamblia*, and parabasalids such as *Trichomonas vaginalis* (27–31).

A second mystery involves the evolutionary dynamics driving transformation of spliceosomal systems, in particular the close association between reduction in intron number and the evolution of homogeneous core splice signals (3). Increased homogeneity of core splice signals implies stronger selection for usage of a particular motif, however why this increased selection should be so closely associated with reduced intron number remains obscure. Two models have been proposed. The ‘selective loss’ model posits increased strictness of spliceosomal requirements evolved first, and has driven loss of introns with sequences not fitting these requirements, leading to a reduced complement of introns strictly adhering to consensus signals (32). The ‘coevolution’ model imagines that intron number reduction occurs first, and that the reduced number of substrates for the spliceosome then allows coevolution of the splicing mechanism and intron sequences (3). To date, no test of the differential predictions of these models have been satisfyingly performed.

A third mystery of intron evolution concerns the gain of spliceosomal introns (33–37). Reconstructions of the history of intron gain over long evolutionary times as well as searches for introns created over more recent times have indicated strikingly divergent evolution. Rates of intron creation show orders of magnitude variation across lineages, with some lineages undergoing little or no intron gain in conserved coding regions over tens of millions of years (38–44) while other lineages experience rapid creation of hundreds or thousands of introns over shorter times (33-37,45). In addition, these recently gained introns appear to have been created by a variety of mechanisms (Curtis and Archibald 2010; Li et al. 2009; Huff et al. 2016; Farlowe et al. 2010, 2011; Hellsten et al. 2011; Worden et al. 2009; Denoeud et al. 2010). Recently, ourselves and others have described intron creation by mobile elements in a variety of lineages (collectively termed Introner Like Elements, or ILEs (33-34,36-37,46-48). Ourselves and others have suggested that ILE dynamics could explain punctate intron gain: most lineages, lacking ILEs, would experience only rare intron gains, occurring by a variety of serendipitous events; by contrast, lineages in which a ILE happens to evolve could undergo rapid intron creation by the genomic spread of these elements (33–34). However, only a small number of lineages undergoing rapid intron creation have been characterized, and characterization of additional cases of intron proliferation is thus crucial.

Here we report investigations of the spliceosomal machinery and introns of *Entamoeba histolytica* and related species. Previous data suggested that *E. histolytica,* a parasite of humans and other primates, is a promising model to study evolution of splicing. *E. histolytica* introns show a strong preference for the atypical 5’ splice site GUUUGU, differing from the typical sequence GUAAGU at an unprecedented two sites (49). Interestingly, previous searches for snRNA sequences in *E. histolytica* identified four of the five core snRNAs, but not U1 (27,49). In addition, initial comparisons between different species of *Entamoeba* suggested substantial intron loss and/or gain of introns (unpublished data associated with (50). Here we report several analyses focused on *Entamoeba* species, which revealed: (i) the presence of a modified U1 snRNA with perfect basepairing potential to the modified 5’ splice site; (ii) a lack of evidence for preferential loss of introns with variant splicing motifs; and (iii) evidence for substantial gain of introns in one lineage of *Entamoeba*.

## Materials and Methods

### Genomic resources

Genome sequences and annotations were downloaded from AmoebaDB 25 for all *Entamoeba* species. Other genomes and annotations were downloaded from Genbank, including *N. vectensis* (assembly ASM20922v1), *Micromonas pusilla* CCMP 1545 (v2.0) and *commoda* (ASM9098v2), *Ostreococcus tauri* (050606) and *lucimarinus* (ASM9206v1), *Bathycoccus* sp TOSAG39-1 (TOSAG39-1), *M. balamuthi* (BN839), *T. trahens* (TheTra_May2010) and *Polysphondylium pallidum* (PolPal_Dec2009). All available Saccharomycotina genomes were downloaded from Genbank on January 15^th^, 2017.

### Intron loss/gain analysis

For all three intron position comparisons (the 5-way *Entamoeba* species comparison, the comparison with *D. discoideum* and *N. vectensis* and the three-way *E.histolytica-E*. *moshkovskii-E.invadens* alignment to identify a maximal set of intron losses), putative ortholog sets were defined by reciprocal blastp searches and were aligned at the protein level in Clustalw with default parameters and then backtranslated with intron sequences added. Conserved regions were defined as those in which each pair of species shared ≥30% protein-level sequence identity over a window of 10 amino acid positions in both directions. For purposes of intron sequence analysis in *Entamoeba*, “confident” intron positions were defined as those that were conserved across all five species. The methods of Roy and Gilbert (17–18) were used to reconstruct intron loss and gain for both datasets (note that while other intron construction methods are available, most methods have been shown to give very similar results in the case of minimal intron gain (20,21); and some of these methods have been argued to be subject to overestimation of parallel intron gains when there are few taxa in the dataset, as here (42). All available ESTs for *M. balamuthi* (20,111 in total) were downloaded from Genbank. A blat search was performed against the *M. balamuthi* genome. Cases where there was a single gap in the alignment between an EST and the genome, and where the gap had GT-AG boundaries, were identified as putative introns.

For analyses of 5’ splice sites in *M. commoda* and *T. trahens*, sequence logos were built from introns containing canonical branchpoint motifs (which are common in these species, e.g., (23)), because previous results have shown a high fraction of false positive intron calls in intron-poor species ((51) and unpublished results). Sequence logos were constructed using Weblogo using default parameters (Crooks et al. 2004).

### Search for U1 candidates in Amoebozoans

Infernal 1.1.2 (including cmsearch) was downloaded from eddylab.org/infernal (53). To search genomes for U1 snRNAs while allowing for sequence flexibility within the basepairing region, the provided U1.cm profile file for cmsearch was modified to give equal scores at these positions for any of the four possible nucleotides. To test this protocol, all U1 snRNA candidate sequences identified by Davila Lopez et al (27) were downloaded from the supplemental materials of that paper. For each of these reported U1 snRNA candidates, the basepairing region was modified in silico to provide Watson-Crick basepairing to GUUUGU (thus the basepairing region was changed from ACUUAC to ACAAAC), and a U1 search was run in cmsearch using the modified U1.cm profile file. In all cases, the real and modified U1 sequences gave identical cmsearch scores, indicating good sensitivity of the method. This method was then run against all downloaded genomes and results parsed using novel Perl scripts.

### Search for U1 snRNA candidates in Saccharomycotina

Among Saccharomycotina species with available genome sequences, 70 gave clear U1 candidates, all with the canonical ACUUAC basepairing sequence. These included *Alloascoidea hylecoeti* (GCA_001600815.1), *Ambrosiozyma kashinagacola* (GCA_001599075.1), *Ambrosiozyma monospora* (GCA_001599995.1), *Ascoidea asiatica* (GCA_001600695.1), *Ascoidea rubescens* DSM 1968 (GCA_001661345.1), *Babjeviella inositovora* NRRL Y-12698 (GCF_001661335.1), *Brettanomyces anomalus* (GCA_001754015.1), *Brettanomyces bruxellensis* CBS 2499 (GCA_000340765.1), *Brettanomyces custersianus* (GCA_001746385.1), *Brettanomyces naardenensis* (GCA_001753995.1), *Candida apicola* (GCA_001005415.1), *Candida arabinofermentans* NRRL YB-2248 (GCA_001661425.1), *Candida auris* (GCF_001189475.1), *Candida boidinii* (GCA_001599335.1), *Candida carpophila* (GCA_001599235.1), *Candida ethanolica* M2 (GCA_001649435.1), *Candida glabrata* CBS 138 (GCF_000002545.3), *Candida homilentoma* (GCA_001599095.1), *Candida infanticola* (GCA_001630515.1), *Candida succiphila* (GCA_001599255.1), *Candida tanzawaensis* NRRL Y-17324 (GCA_001661415.1), *Candida tenuis* ATCC 10573 (GCF_000223465.1), *Candida versatilis* (GCA_001600375.1), *Cyberlindnera fabianii* (GCA_001599195.1), *Cyberlindnera jadinii* NBRC 0988 (GCA_000328385.1), *Debaryomyces fabryi* (GCF_001447935.1), *Debaryomyces hansenii* CBS767 (GCF_000006445.2), *Eremothecium coryli* CBS 5749 (GCA_000710315.1), *Geotrichum candidum* (GCA_000743665.1), *Hyphopichia burtonii* NRRL Y-1933 (GCA_001661395.1), *Kazachstania naganishii* CBS 8797 (GCA_000348985.1), *Komagataella pastoris* (GCA_001708105.1), *Komagataella phaffii* GS115 (GCF_000027005.1), *Komagataella phaffii* GS115 (GCF_000027005.1), *Kuraishia capsulata* CBS 1993 (GCA_000576695.1), *Lipomyces starkeyi* NRRL Y-11557 (GCA_001661325.1), *Meyerozyma caribbica* MG20W (GCA_000755205.1), *Meyerozyma guilliermondii* ATCC 6260 (GCF_000149425.1), *Millerozyma acaciae* (GCA_001600675.1), *Millerozyma farinosa* CBS 7064 (GCF_000315895.1), *Nadsonia fulvescens* var*. elongata* DSM 6958 (GCA_001661315.1), *Nakazawaea peltata* (GCA_001599355.1), *Ogataea parapolymorpha* DL-1 (GCF_000187245.1), *Ogataea polymorpha* (GCF_001664045.1), *Pachysolen tannophilus* NRRL Y-2460 (GCA_001661245.1), *Pichia kudriavzevii* (GCA_000764455.1), *Pichia membranifaciens* NRRL Y-2026 (GCF_001661235.1), *Priceomyces haplophilus* (GCA_001599895.1), *Saccharomyces pastorianus* CBS 1513 (GCA_000586595.1), *Saccharomycopsis malanga* (GCA_001599215.1), *Scheffersomyces stipitis* CBS 6054 (GCF_000209165.1), *Spathaspora arborariae* UFMG-19.1A (GCA_000497715.1), *Spathaspora girioi* (GCA_001657455.1), *Spathaspora gorwiae* (GCA_001655765.1), *Spathaspora hagerdaliae* (GCA_001655755.1), *Spathaspora passalidarum* NRRL Y-27907 (GCF_000223485.1), *Sporopachydermia quercuum* (GCA_001599295.1), *Starmerella bombicola* (GCA_001599315.1), *Sugiyamaella lignohabitans* (GCF_001640025.1), *Tortispora caseinolytica* NRRL Y-17796 (GCA_001661475.1), *Wickerhamia fluorescens* (GCA_001599155.1), *Wickerhamiella domercqiae* (GCA_001599275.1), *Wickerhamomyces anomalus* NRRL Y-366-8 (GCF_001661255.1), *Wickerhamomyces ciferrii* (GCF_000313485.1), *Yarrowia deformans* (GCA_001600075.1), *Yarrowia keelungensis* (GCA_001600195.1), *Yarrowia lipolytica* CLIB122 (GCF_000002525.2), *Yarrowia* sp. JCM 30694 (GCA_001600515.1), *Yarrowia* sp. JCM 30695 (GCA_001602355.1), and *Yarrowia* sp. JCM 30696 (GCA_001600535.1).

### Branchpoint analysis

To find branchpoints, branchpoints for 100 introns for each species were identified by hand, and a position weight matrix (PWM) built for these branchpoints, and the fraction of scored branchpoints falling at various positions relative to the 3’ splice site tallied. These matrices were then used to provide a score for each adenine nucleotide within the last 30 nts of each intron, and the adenine with the largest score chosen as the predicted branchpoint. This approach was used to predict branchpoints for an additional 100 introns, which were also predicted manually.

The automated and manual predictions corresponded well for *E. histolytica, E. moshkovskii* and *E. invadens* (greater variation in branchpoint position in the other *Entamoeba* species, coupled with the high rate of branch-point-like motifs in these AT-rich species (e.g., ATTAAT), thwarted confident determination of branchpoints in these two species. For Figure 6, branchpoint scores were normalized to a value from 0 to 1, where 1 represents the maximum possible score (for both *E. histolytica* and *E. moshkovskii*, this was for the motif ATTA<u>A</u>T, where the underline indicates the branchpoint A) and 0 the minimum possible score (for both species, for the motif CGGGC<u>G</u>C). Randomization tests were run by novel Perl scripts by selecting random subsets of the entire set of introns and comparing this to the real observed subset.

### Spliceosomal protein searches

Spliceosomal protein searches were performed on proteome assemblies available from NCBI and UniProt. A list of relevant human spliceosomal proteins was used as queries in local BLASTp (version 2.9.0+) searches against independent proteome databases (initial e-value threshold of 10^-6^) (54). The results from the BLAST searches were further screened by analyzing domain content (HMMsearch, HMMer 3.1b2 – default parameters), size comparisons against human protein sequence length (within 25% variation), and reciprocal best-hit BLAST searches (RBH) to the query proteome (55–57). To avoid bias in protein domain content, domains used for HMM searches were defined as in Hudson et al. (26). Briefly, a conserved set of domains for each spliceosomal protein was assembled by using only those domains present in all three of the human, yeast, and *Arabidopsis* orthologs. Fungal ortholog candidates in this study were scored and awarded a confidence value of 0-9 based on passing the above criteria. Scores were calculated by starting at 9 and penalizing candidates for falling outside of the expected size range (−1 point), missing HMM domain calls (−2 points), and failing to strictly pass RBH (−5 points). A more “relaxed” RBH protocol was used and proteins that found reciprocal hits in the top three candidates started at a score of 4 and were reduced due to HMM and size penalties as above. Finally, ortholog candidates that only had initial BLAST hits but failed all other tests were assigned a score of 0.5 to differentiate them from queries with no BLAST results. Only ortholog candidates receiving a score of 9 were used for spliceosome protein counts and percentage calculations. The curated subset of the protein list is representative of highly-studied protein factors.

Presence/absence scores for each protein in each species are shown in Supplemental Table 1.

## Results

### Atypical donor splice sites and constrained branchpoint structures are common features of Entamoeba species

A previous study reported that *Entamoeba histolytica* shows an atypical donor splice site in which both +3 and +4 sites show atypical nucleotides (GUUUGU, as opposed to the GUAAGU consensus most commonly found among eukaryotes; Davis et al. 2007). To explore this pattern further, we downloaded full genomic sequences for all 5 *Entamoeba* species for which genomes have been sequenced and studied donor splice sites at confident intron positions (Figure 1a,b; see Materials and Methods). All five species exhibited the atypical GUUUGU splice site previously reported for *E. histolytica*. Between 87.9 and 89.5% of introns exhibit a GUUUGU donor site and between 96.9% and 98.5% exhibit GUUUGN, making *Entamoeba* among the eukaryotes with the strictest splicing signals. In addition, all 5 species exhibited a strong preference for a U at the 7^th^ position (77.7-83.1% of introns across species exhibiting a 7U), a position for which most characterized species do not show a strong preference (Figure 1a (9)). Thus all characterized species within the genus *Entamoeba* share a preference for an atypical GUUUGUU donor splice site. Notably, this atypical GUUUGU motif could potentially allow for increased basepairing with the U6 snRNA’s basepairing region, in which the sequence 5’-ACAGA-3’ basepairs with positions 2-6 of the 5’ splice site (i.e., G<u>UUUGU</u> in *Entamoeba*; note that *Entamoeba* retains the typical ACAGA U6 motif (27)).

**Figure 1.**
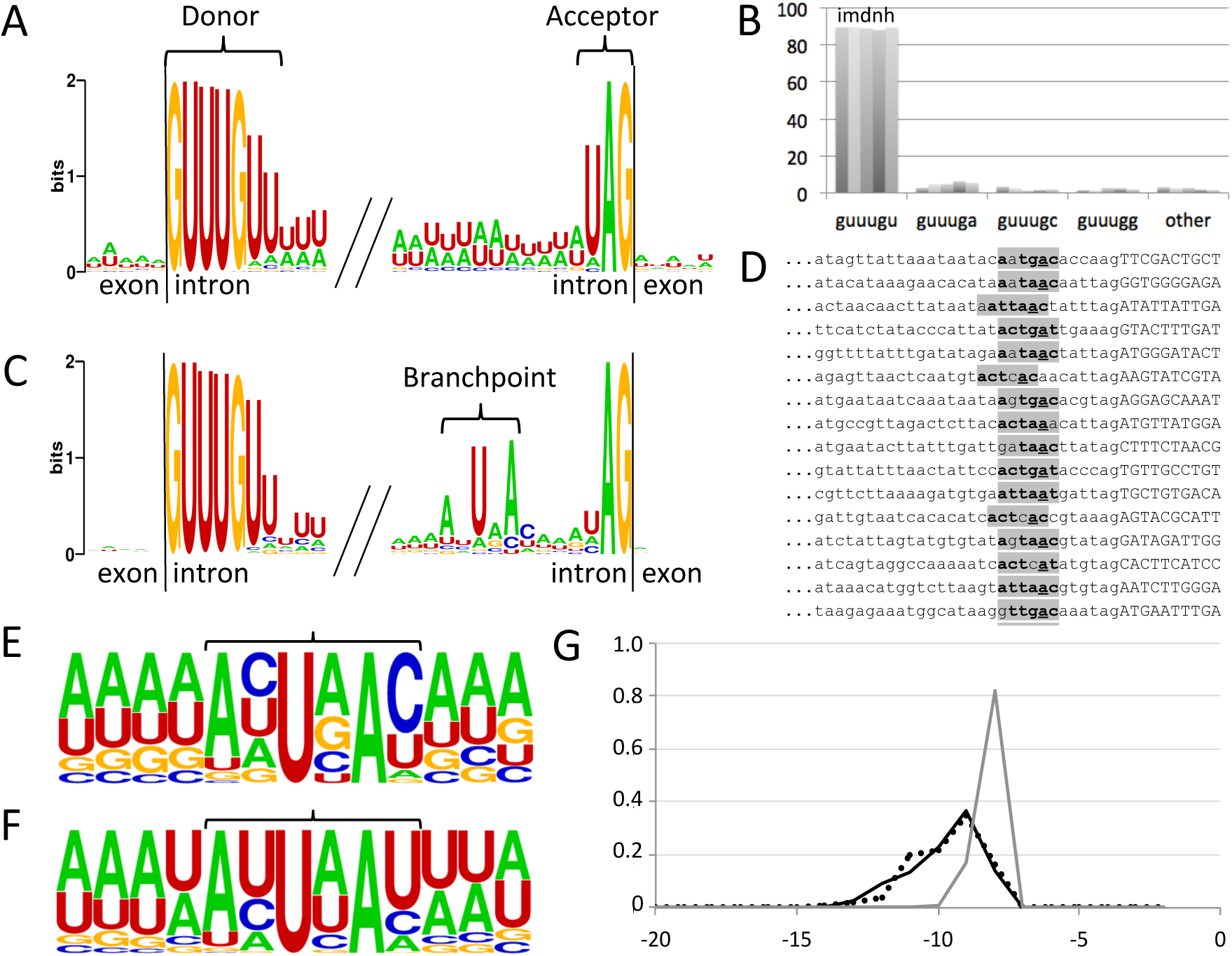
Core splicing motifs in *Entamoeba* species show a high degree of homogeneity. A. Introns in *E. histolytica* show a highly conserved atypical extended donor splice site, with atypical usage of uracil at the 3^rd^ and 4^th^ intronic position and clear preference for uracil at the 7^th^ intronic position as well. B. Percentage of all confident introns with various splice boundaries for five species of *Entamoeba*, showing a clear preference for all species (*i/m/d/n/h* indicates *E. invadens/moshkovskii/dispar/nuttalli/histolytica*). C. Splicing motifs for *E. invadens*, showing a strong preference for branchpoint position eight nucleotides before the 3’ splice site. D. 3’ sequences for a randomly chosen subset of *E. invadens* introns, showing sequences and positions of branchpoints. E.,F. Branchpoint motifs (bracket) and surrounding sequence for predicted branchpoints in *E. invadens* (E) and *E. histolytica* (F). G. Relative positions for predicted branchpoints relative to the 3’ splice site for *E. histolytica* (dotted line), *E. moshkovskii* (black line) and *E. invadens* (gray line).

Interestingly, the sequence logo for the most distantly-related of the five species, *E. invadens*, showed clear nucleotide preference for a TNA motif near the beginning 10 nucleotides upstream of the 3’ splice site (Figure 1c). Scrutiny of individual sequences revealed a clear AYTRAY pattern, which is the canonical branch point sequence (Figure 1d). Introns that did not contain a candidate motif at the exact site typically revealed a putative branchpoint motif at a nearby site (almost always one or two nucleotides upstream (Figure 1d)). Out of 904 *E. invadens* introns at positions of conserved protein alignment, 98% had a TNA either 10 nts (81.2%) or 11 nts (16.8%) away from the 3’ splice site, and 16/18 remaining introns had a TNA either 9 or 12 nts away. The putative branchpoint motif matched the consensus AYURAY at 5 or 6 out of 6 positions for 87.6% of introns. Alignment of these motifs (both at the most common and neighboring sites) reveals the overall branchpoint motif shown in Figure 1e. Thus, *E. invadens* shows a strong conservation of the branchpoint position along the intronic sequence, as previous observed in several other highly transformed spliceosomal systems including diplomonads, parabasalids and some yeasts within Saccharomycotina (2,23–24,58–59). Scrutiny of 3’ regions of intronic sequences from other *Entamoeba* species also revealed candidate branchpoint motifs for large numbers of introns, albeit with more positional flexibility (e.g., Figure 1f). Notably, whereas the clear preference for a branchpoint position of -8 nts in *E. invadens* makes identification of putative branchpoint motifs straightforward, the greater flexibility in other *Entamoeba* species coupled to the low sequence complexity of both the motif (many putative branchpoint motifs are AUUAAU) as well as the intron sequences in general (76-83% A/U) means that branchpoint-like motifs occur by chance frequently in an intron, complicating confident identification of a single candidate branchpoint per intron.

### Development of a modified search protocol to search for atypical U1 snRNA genes

In order to search for U1 snRNAs while allowing for the possibility of atypical basepairing regions, we used Infernal (53) to search genomes for candidate U1 sequences using a modified profile file to allow for flexibility specifically at the positions of the U1 snRNA that basepair with the +3 and +4 positions of the intron (see Materials and Methods). In order to test the sensitivity of this method, we generated a test file containing both the observed ‘standard’ U1 snRNAs reported for a wide variety of species (all sequences reported in (27)) as well as variants of these sequences modified in silico to provide a variety of sequences within the basepairing region. Running the modified protocol on these sequences showed identical sensitivity to the standard and modified snRNAs, giving identical scores for the corresponding standard/modified pairs, indicating success of the method in allowing for a diversity of sequences at the basepairing site (data not shown).

### Identification of an atypical U1 snRNA in Entamoeba

We used this modified pipeline to search for candidate U1 snRNA genes in the genomes of five *Entamoeba* species. For each species this search revealed a single sequence exhibiting the core characteristics of a U1 snRNA, including a secondary structure consisting of four stem-loop (SL) structures, with putative binding sites for the 70k and U1a proteins in SL I and II, respectively, and a strong candidate SM binding site between SLs III and IV (Figure 2a,b). The candidate U1 snRNA genes for each of the five species show a modification at the basepairing site, with the sequence ACAAAC observed in place of the canonical ACUUAC. Notably, this sequence provides exact Watson-Crick basepairing potential with the modified *Entamoeba* donor site GUUUGU (Figure 2a,b). In all five species, the preceding base is also an ‘A’ (AACAAAC), potentially allowing for perfect basepairing with the extended donor site GUUUGUU. Interestingly, significant differences in the structures and lengths of SL III and IV were observed between species, with *E. moshkovskii* showing transformed sequences relative to the other four species (insets in Figure 2a). At the deepest divergences within the genus, even more substantial differences were observed, with *E. invadens* U1 sequence differing from that of the other species at a large number of sites. For instance, 6/20 bases in the stem of SL I are different, with all changes maintaining basepairing potential, and 17/31 sites in the stem of SL II are different, including differences involving the identities of pairing partners (grey basepairs in Figure 2b). Notably, for all five species, the upstream sequence is highly A-rich (Figure 2c), as is found for other spliceosomal snRNAs in this genus (Figure 2d), a common feature of Pol III promoters.

**Figure 2.**
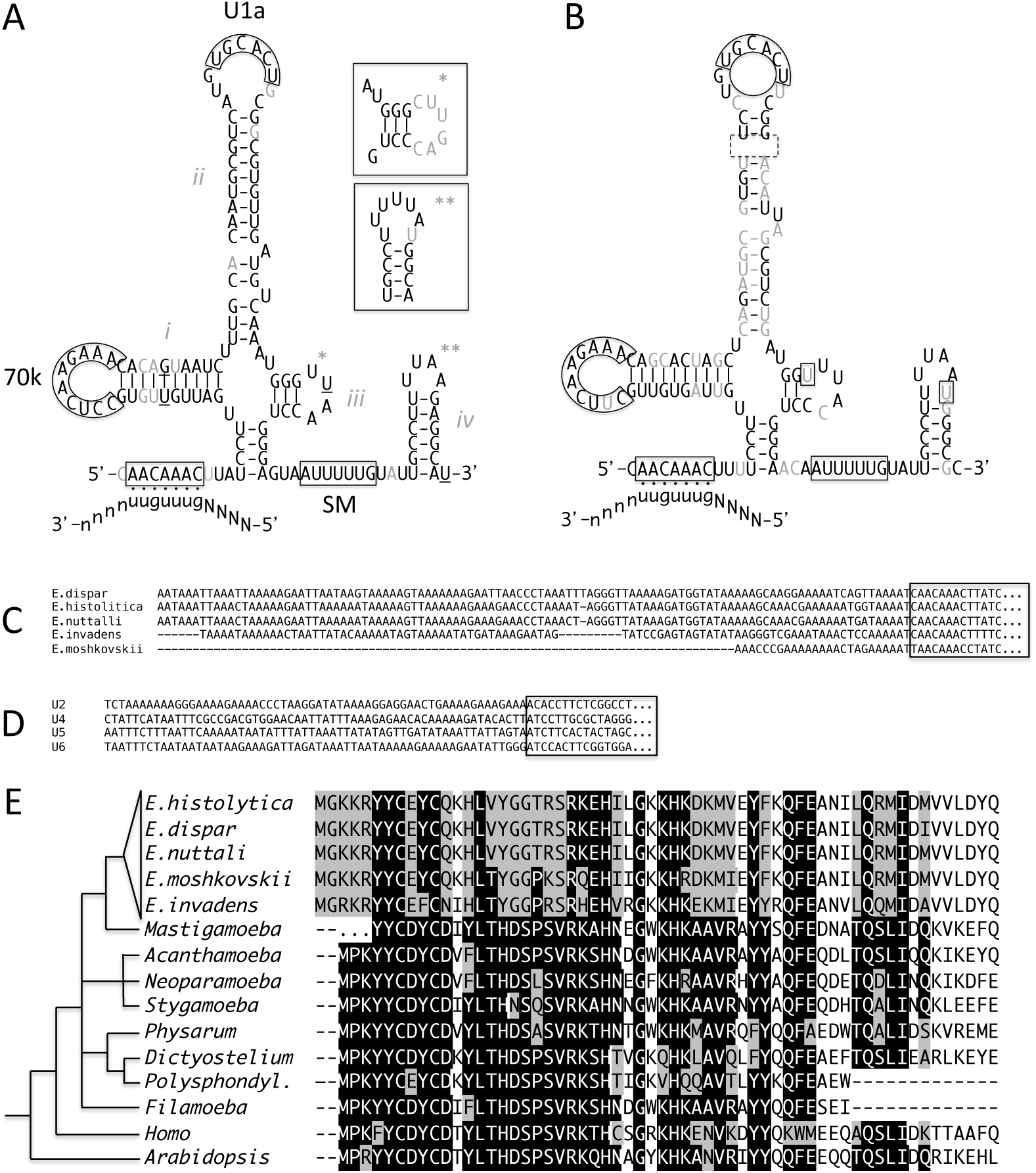
Characteristics and contexts of U1 snRNP in *Entamoeba.* A. U1 snRNA sequence and predicted secondary structures for *Entamoeba* species. Sequence shown represents the consensus sequence between the closely-related species *E. dispar, E. nuttalli* and *E. histolytica*. Boxed sequences represent binding sites for the 5’ splice site (bottom left), 70k, U1a and SM proteins, as indicated. Labels *i*-*iv* indicate stem-loops 1-4. Underlined nucleotides represent sites at which a transition difference is observed between *E. dispar/nuttalli/histolytica* (no transversions are observed). Grey nucleotides indicate sites at which a transition difference is different between the consensus sequence and *E. moshkovskii* sequence (no transversions are observed other than in regions indicated in inset boxes). Inset boxes indicate variant SL III and SL IV sequences observed in *E. moshkovskii*. B. U1 snRNA sequence for *E. invadens*. Basepair substitutions relative to *E. dispar/nuttalli/histolytica* consensus (part A) are shown in gray. Dotted box shows deletion of a basepair relative to part A. Gray basepairs indicated basepairings between nonhomologous pairs relative to part A. Boxed single nucleotides represent positions predicted to undergo basepairing in part A but not in part B. C. Alignment of genomic regions upstream of U1 snRNA genes in *Entamoeba* species, showing extended A-rich regions in all five species. The box indicates the beginning of the predicted U1 sequence. D. A-rich regions upstream of other predicted spliceosomal snRNA genes in *Entamoeba.* Box indicates the beginning of the predicted snRNA sequences. Sequences shown are from *E. invadens*. E. A highly-diverged U1c candidate in *Entamoeba*. An alignment of the highly-conserved *N* terminal region of U1c from five *Entamoeba* species, the relative *Mastigamoeba balamuthi*, seven distantly related amoebozoa species, human and *Arabdiopsis*, showing that *Entamoeba* shows a greater degree of deviation (grey boxes) and lower degree of adherence (black boxes) at strongly conserved amino acid positions (non-boxed positions indicate evolutionarily variable amino acid sites).

These atypical motifs in the U1 snRNA and 5’ splice sites suggested the possibility of an altered mechanism for 5’ splice site recognition in *Entamoeba.* Interestingly, BLASTP searches of core U1 snRNP proteins from humans against *Entamoeba* species identified candidates for most proteins (data not shown), but did not identify a candidate for U1c, the protein responsible for 5’ splice site recognition in studied eukaryotes (60). Further searches relying on homologs from other species from within amoebozoans identified a potential highly diverged U1c candidates, which is much shorter than most known U1c proteins (96 amino acids compared to >150 for most species), and which shows a number of changes at conserved amino acid sites since the divergence from the related species *Mastigamoeba balamuthi* (Figure 2e). For instance, the observed sequences require a minimum of 39 changes at highly-conserved sites within this single genus, compared to 25 across the rest of surveyed amoebozoans (and none within *M. balamuthi* over the same time.

Interestingly the blast searches did identify a clear candidate for NAM8/TIA-1 (data not shown), a protein involved in splicing of splicing of introns with nonconsensus 5’ splice sites, which might have been expected to be dispensable in organisms with such strict adherence to the 5’ splice site consensus 5’ motif.

### Complementary U1 snRNA and donor site a related intron-rich amoeba with a different atypical donor splice site

To trace the evolutionary history of the *Entamoeba* U1 snRNA, we also searched for candidate U1s in the genome of *Mastigamoeba balamuthi*, the one non-*Entamoeba* species within the large group Archamoeba for which a genome is available. Available gene structures suggest a much higher intron density (2.5 introns per gene), suggesting that massive intron loss occurred in the ancestor of *Entamoeba* following divergence from *Mastigamoeba*. Again allowing for flexibility at the basepairing sites, this search revealed a clear U1 candidate in this species. Again, this species exhibited classical hallmarks of a U1 snRNA (Figure 3a). However, this sequence showed neither the classical sequence nor the *Entamoeba* variant but a third variant, ACGUAC, which would provide Watson-Crick basepairing with GUACGU.

**Figure 3.**
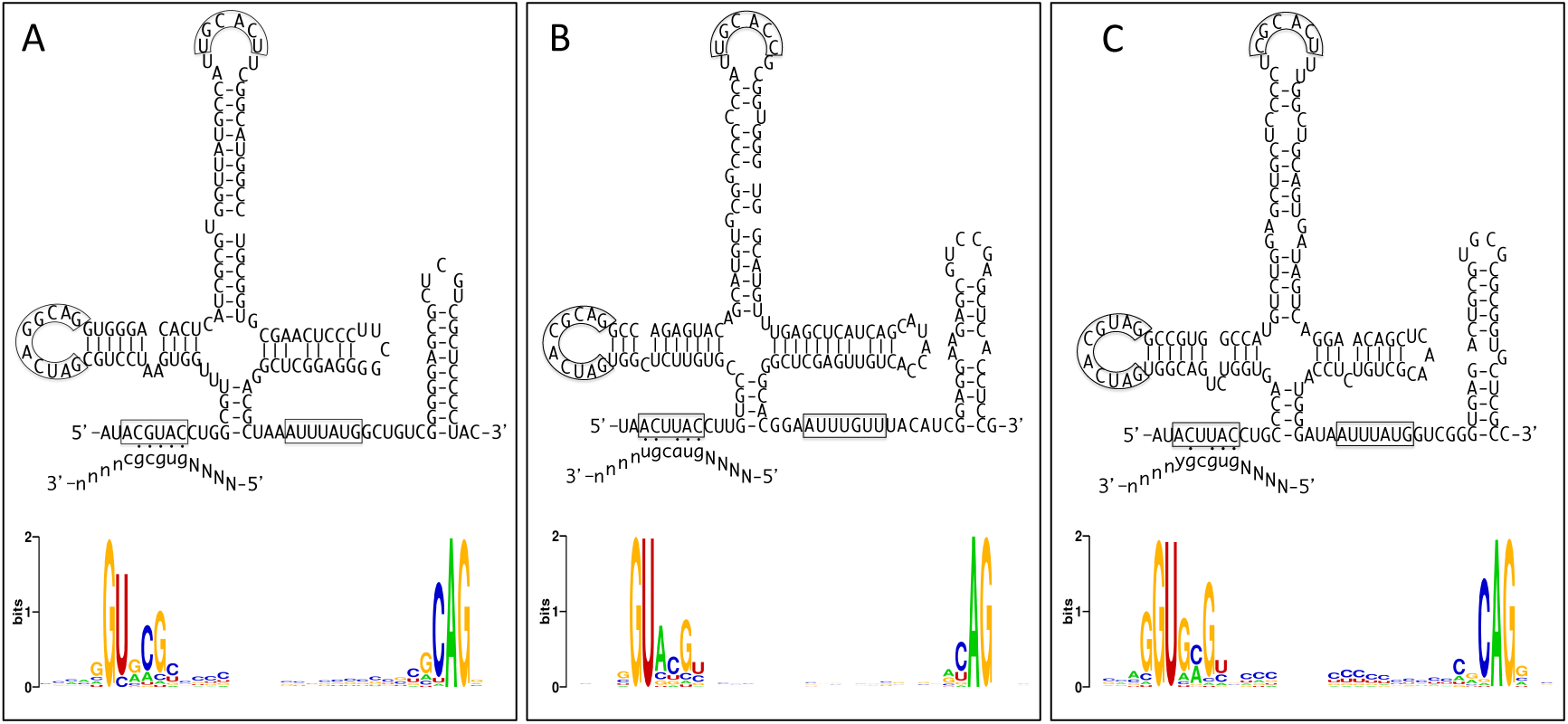
Splice site motifs and predicted U1 snRNA sequences and structures for three species with a preference for a pyrimidine (C/U) at the fourth intronic position. Compensatory change between U1 snRNA and splice site is observed for the amoebozoan *M. balamuthi* (A), but not for the prasinophyte *M. pusilla* CCMP1545 (B) or the apusomonad *T. trahens* (C). Splice site motifs for *M. pusilla* exclude introns from the AT-rich atypical chromosome (32).

Genome-wide intron-exon structures of *M. balamuthi* are not yet available to our knowledge. However, blat searches of available EST sequences from this species revealed a number of splicing events. Interestingly, donor sites for the identified introns showed a clear preference for a different sequence GUGCGC (Figure 3a). This sequence provides Watson-Crick basepairing for the modified site in the U1 snRNA (4C basepairing with the atypical U1 ‘G’), as well as wobble basepairing at the neighboring site (3G pairing with the standard U in the U1 snRNA). Notably, the atypical 6C of donor splice sites does not provide canonical basepairing potential with the typical A nucleotide found at the potential pairing site for either the standard U1 sequence or for the *M. balamuthi* U1 site. In addition, unlike *Entamoeba*, no potential for extended basepairing is observed (Figure 3a).

### Search for U1 snRNAs in species with atypical donor splice sites

Including *Entamoeba*, there are six known lineages in which intron-exon structures have independently been transformed by both massive intron loss and evolution of a preferred donor site that does not provide standard basepairing with the classic U1 snRNA sequence. U1 snRNAs have been reported for representatives of four of these lineages: *Giardia intestinalis* (a diplomonad), *Trichomonas vaginalis* (a parabasalid) and *S. cerevisiae* and some relatives (saccharomycotina), and now *Entamoeba* (28–31).

To characterize U1 snRNAs in these intriguing lineages, we used the modified protocol described above to search for U1 snRNA genes. First, we searched for a U1 snRNA sequence in the genome of *Thecamonas trahens*, a representative of the poorly-studied eukaryotic group Apusozoa, which exhibits a preference for C at the 4^th^ intronic position (Figure 3b). This search revealed a single strong U1 snRNA candidate with the standard basepairing sequence, indicating lack of a complementary change in the U1 (Figure 3b). Similarly, we performed searches against five available representatives of Mammieles, a group of green algae which exhibit a preference for GYGCGY donor sites across most of their genome (Figure 3c; although the presence of very different splicing signals for one atypical chromosome in this species complicates this classification (61)). Again, for all five species for which a candidate was identified, the single strong candidate exhibited the standard basepairing region, indicating lack of compensatory changes in the U1 snRNA. Figure 3c shows the obtained sequence for one of the five, *Micromonas pusilla* CCMP1545.

Saccharomycotina yeasts are a group which includes the baker’s yeast *S. cerevisiae*, many of which have a clear preference for +4U in their donor sites. This group includes more than 100 species with available genomes, allowing an unprecedented opportunity to search for transformed U1 snRNAs. Searches of these genomes using the modified protocol yielded 69 species with candidate U1 snRNA sequences. Notably, all 69 species show U1 snRNA candidates with the standard pairing (as in *S. cerevisiae*); none showed a candidate with specifically modified basepairing region to provide standard basepairing with the modified GUAUGU donor site (the list of species is given in Materials and Methods).

### Characterization of spliceosomal proteomes in Entamoeba and other Amoebozoans

Given the surprising transformation of a core snRNA and previous evidence for an association between intron density and the complexity of the spliceosome, we next sought to characterize the spliceosomal machinery across *Entamoeba*. We combined information on ancient spliceosomal proteins with factors known to associate with the *E. histolytica* spliceosome from a previous mass spectrometry study to compile a list of putative spliceosomal factors. We used sensitive methods to detect homologs of these proteins in each genome in order to characterize the complement of spliceosomal proteins in *Entamoeba* and other Amoebozoans.

The five *Entamoeba* species in this study showed a markedly reduced set of spliceosomal proteins (Figure 4, Supplemental Table 1). Reductions in protein counts occurred in all particles. Interestingly, the smallest amount of change was recorded in most core snRNP-associated protein classes (namely U2, U4/U6, U4/U6.U5 tri-snRNP, and U5 structures). Interestingly, U1 proteins did not show particular conservation; however it is difficult to ascertain whether this reflects faster evolution of U1 proteins due to changes in U1 function, generally faster evolution of this non-catalytic snRNP, or simply stochastic fluctuation given the generally small number of U1 proteins. The patterns of *Entamoeba* protein losses are shared between the species with no clear differences between the organisms. These patterns, however, when solely looking at protein conservation/ loss, do mirror patterns observed in the reduced spliceosome of *Saccharomyces cerevisiae*.

**Figure 4.**
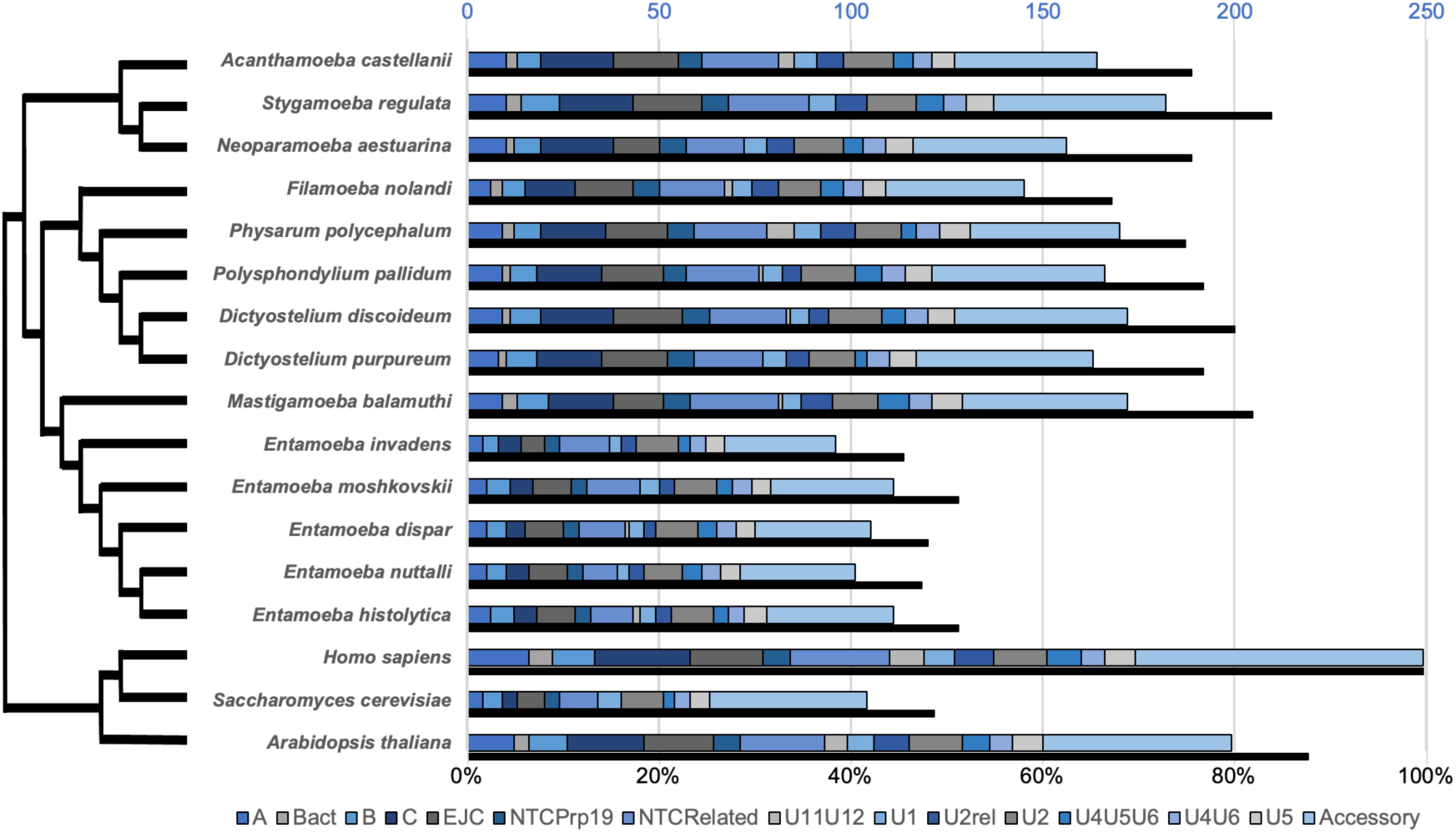
Reduction of the spliceosome in *Entamoeba*. The lower solid bar represents the percentage of select human spliceosomal proteins with putative orthologs detected in each species, for a curated set of proteins (see Methods and Supplemental Table 1). The upper segmented bars represent the number of all human spliceosomal proteins with putative orthologs detected in each species per spliceosomal group. “Accessory” proteins include cap-binding proteins (CBP), disassembly proteins, hnRNP, LSm, misc, mRNA, mRNP, RES, Sm, SR, and Step2Protein groups. Phylogeny based on Kang et al. (78).

### Ongoing intron loss and lineage-specific intron gain in Entamoeba

In order to better understand the history of splicing in *Entamoeba*, we sought to study intron loss and gain among the five species within the genus with available genomes. Putative orthologs were identified between the five species and available outgroup species and standard methods used to reconstruct intron loss and gain in the history of the genus (Figure 5a (17–18)). This produced a dataset of 1025 unique intron positions within 2344.1 kb of conserved coding regions in 3043 sets of orthologs. Within ingroups, this reconstruction showed a clear excess of intron losses over gains, as has previously been found for various other groups of eukaryotes (17-18,38-44,62-63). Interestingly, this analysis also revealed a large number of introns that were specific to the most distantly related species, *E. invadens* (Figure 5a).

**Figure 5.**
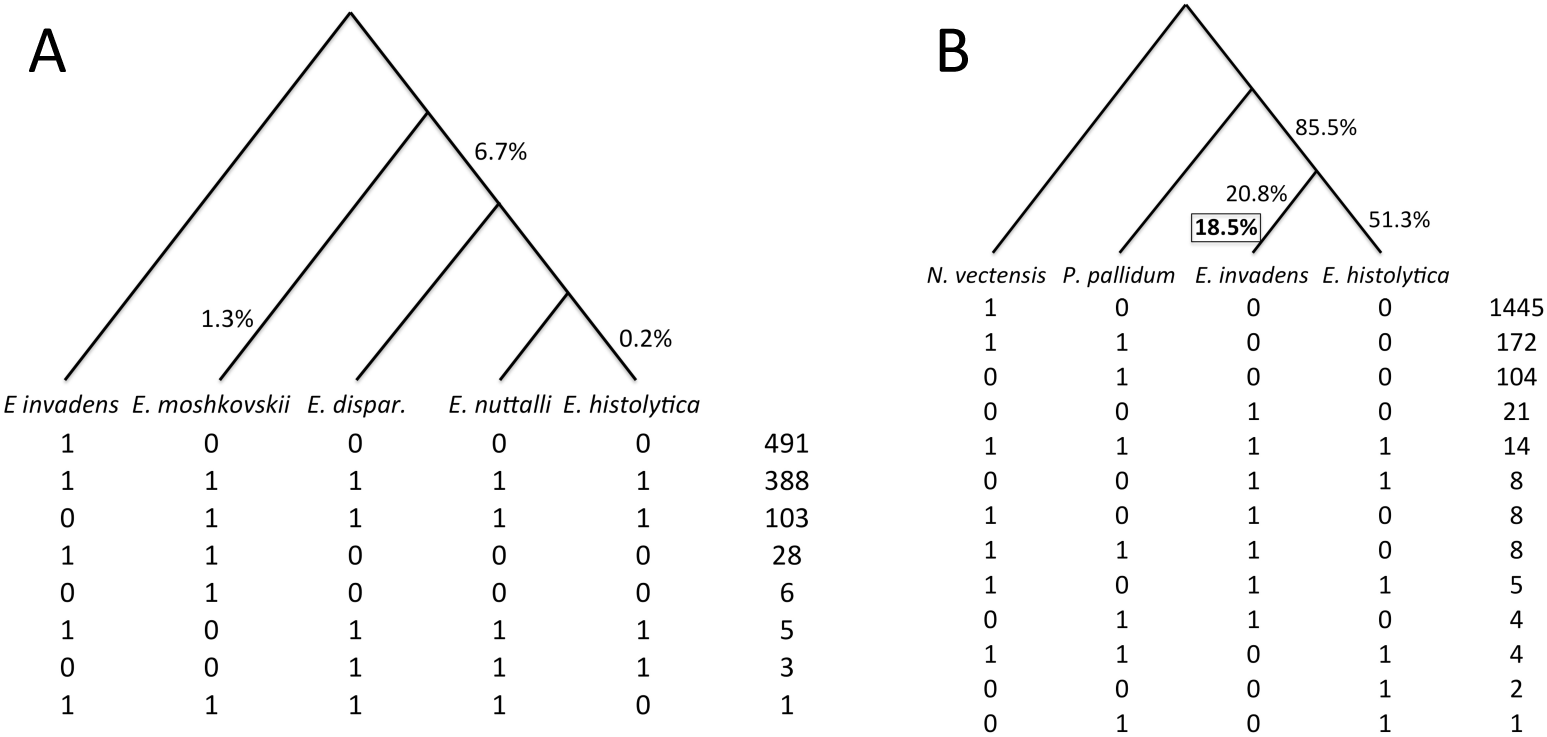
Reconstruction of intron loss and gain in conserved coding regions in the evolutionary history of *Entamoeba,* within five species within the genus (A) and over longer time scales (B). Below each tree, the numbers of intron positions in conserved coding regions showing presence/absence (1/0) are shown. Unboxed values on the tree show the fraction of introns estimated to have been present in the ancestral node that are estimated to have been lost along the corresponding internal/external branch. Estimated degree of loss along branches without an indicated value is not significantly different from zero. Boxed value in part B indicates the fraction of introns in *E. invadens* estimated to have been gained since the divergence with *E. histolytica*.

In order to determine whether these *E. invadens-*specific intron positions reflected intron loss or intron gain, we then performed way comparisons between *E. invadens*, *E. histolytica* and two outgroup species, the Amoebozoan slime mold *Polysphondylium pallidum* and the animal *Nematostella vectensis* (chosen because of its highly ancestral complement of intron positions (64)). This produced a dataset of 1796 unique intron positions (including 75 in one or both *Entamoeba* species) within 188.2 kb of conserved coding regions in 598 sets of orthologs. These comparisons revealed that, whereas many of the intron positions shared across the genus are also represented in outgroups, introns found in *E. invadens* but not other *Entamoeba* species were not represented in outgroups (Figure 5b). This is just as expected if the *E. invadens-*specific introns represent intron gains in that lineage (rather than losses in other *Entamoeba* species). We used a previous method to reconstruct intron loss and gain in the history of *Entamoeba* (17–18), revealing substantial loss in various branches within the genus as well as in the ancestor of the genus, as well as substantial intron gain in *E. invadens* (Figure 5b).

### No evidence for preferential loss of introns with suboptimal splicing signals

We next sought to test whether introns with putatively suboptimal splicing signals were more likely to be lost, as previously proposed (32). To maximize the number of observed changes, we generated a dataset of intron positions at conserved coding positions, including 1623 unique intron positions within 3432.6 kb of conserved coding regions in 4433 ortholog sets. This included 37 putative intron losses in *E. histolytica* (those present in *E. moshkovskii* and *E. invadens*), 10 putative intron losses in *E. moshkovskii* (present in *E. histolytica* and *E. invadens)*, and 187 losses in *E. invadens* (those present in *E. histolytica* and *E. moshkovskii*, which are likely to be losses in *E. invadens*, given the lack of evidence for intron gain in the *E. histolytica/E. moshkovskii* branch(es) of the tree (Figure 5b). To compare introns lost and retained in *E. histolytica*, we used sequences of introns at the homologous position in *E. moshkovskii*, and vice versa. For *E. invadens* we used sequences from *E. histolytica*. For each comparison, lost introns were compared with 100,000 subsets of introns equal size generated from random sampling from all introns (lost plus retained) for that branch (see Methods for details).

First, we compared the fraction of lost introns exhibiting the putatively optimal 5’ splice site with random subsets. For instance, for 29 out of the 37 introns lost in *E. histolytica*, the *E. moshkovskii* intron position at the homologous position contained the full GUUUGUU 5’ splice site. Among 100,000 random subsets of 37 introns drawn from all 689 *E. moshkovskii* introns in the set (those either retained and lost in *E. histolytica*), 77786 had at least 29 or fewer introns with the full splice site motif, thus for the test of whether introns with suboptimal extended 5’ splice site motifs are more likely to be lost, *P* = 0.78 (top, Figure 6a). The analogous test was performed for the core GUUUGU splice site (*P* = 0.90, top, Figure 6b). These tests were performed for all three lineages as well as for the total across lineages (Figure 6a,b). For 6/8 comparisons, lost introns had a (non-significantly) larger fraction of introns with the putative optimal splice site, opposite the prediction of preferential loss of suboptimal introns. In no case was the comparison statistically significant in either direction. Second, to compare the average adherence to the branchpoint motif (as evaluated by a PWM branchpoint score, see Methods) for lost and retained introns, we compared the average branchpoint score for lost introns with 100,000 random subsets of all introns (Figure 6c). None of the four comparisons were close to statistically significant, with *P*-values ranging from 0.22-0.58. Thus we found no evidence for preferential loss of introns with suboptimal splicing motifs.

**Figure 6.**
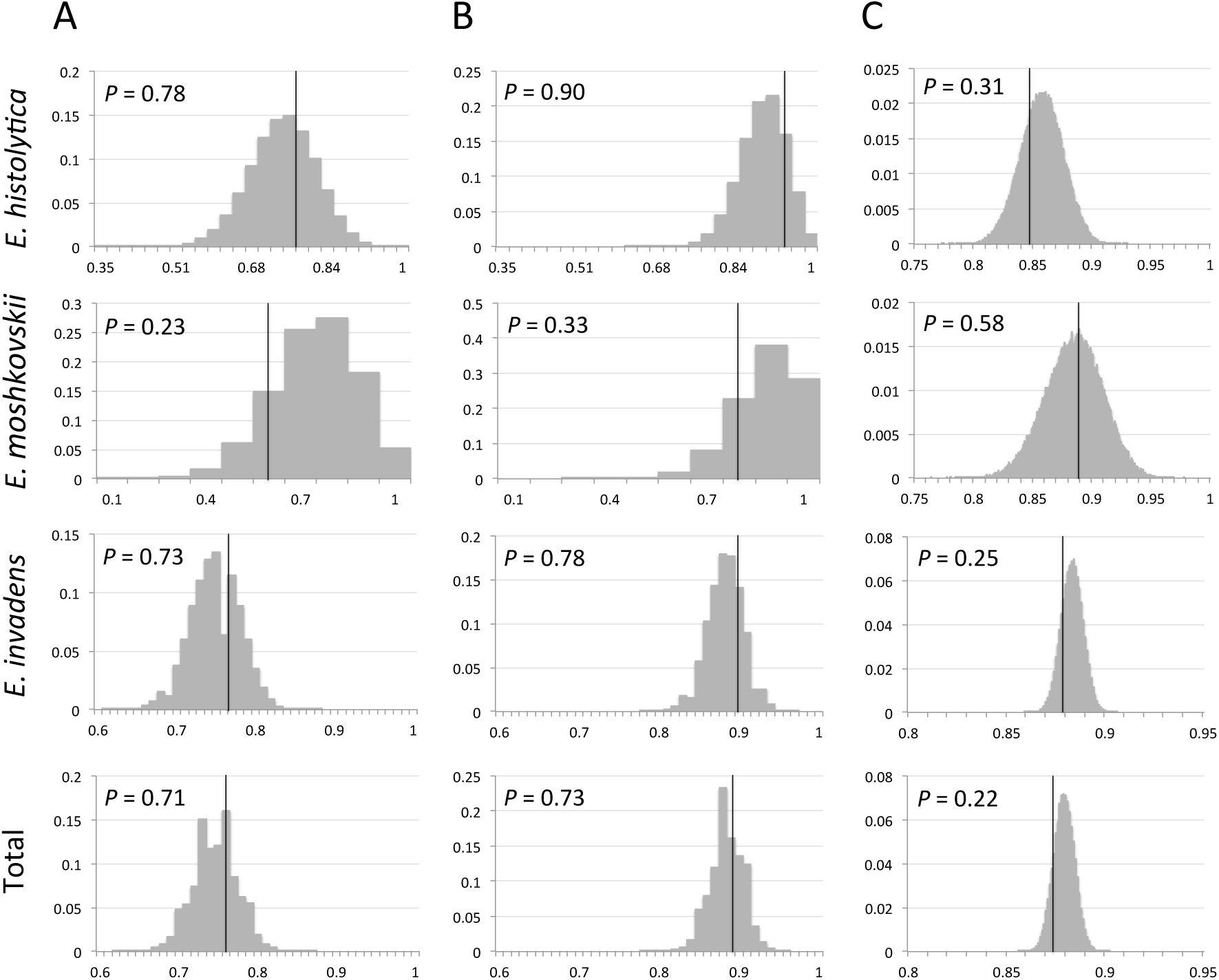
Introns with stronger splice sequence motifs are not more likely to be lost. Comparison of real (vertical black bar) sets of lost introns with 100,000 randomly chosen subsets (grey histogram) for three species individually and the totals across species. A,B. Proportion of introns with optimal 5’ splice site for extended GUUUGUU motif (A) or core GUUUGU motif (B). C. Distribution of average branchpoint score.

### Gained introns in E. invadens show strong similarity of splicing signals to ancestral introns

We next searched putative intron gains in *E. invadens* for the presence of distinctive signatures predicted by different reported and proposed mechanisms of intron gain (see Discussion). *E. invadens* introns in conserved coding regions were divided into potential intron gains and likely ancestral introns based on whether introns were found at that position in other *Entamoeba* species. We first tested whether there was evidence for clearly recognizable sequence homology between introns by performing BLASTN searches between potential intron gains. This BLAST search returned no promising candidates for between-intron homology. We next sought evidence for greater similarity between the regions spanning the 5’ and 3’ intron boundaries for potential intron gains, as is predicted by intron gain mechanisms that involve formation of staggered double stranded breaks (35,62). We did slightly greater similarity between 5’ and 3’ boundaries for potential intron gains, however this was entirely explained by potential intron gains having a greater tendency to have the optimal guanine nucleotide at the exonic bases directly before (30.7% v. 18.6%; *P* < 0.01 by a chi-square test) and after (29.7% v. 19.5; *P* < 0.01) the intron. This greater adherence to the optimal splicing context for newly-gained introns has been found previously (65–66). Finally, we tested the degree of adherence of the potential intron gains to the general consensus sequence. Putative intron gains were not more likely to exhibit the core splice site GUUUGU (92.1 v 91.7%; *P* > 0.1 by a chi-square test), but were more likely to exhibit the extended GUUUGUU splice sites (73.3% v 66.4%; *P* = 0.03 by a chi-square test). No difference was found in the fraction of introns with a branchpoint at the preferred -8 position (80.3% for putative gains versus 81.9% for putative ancestral introns, *P* > 0.1 by a chi-square test), nor in the adherence to branchpoint score, as evaluated by a PWM approach (*P* > 0.1 by randomization).

## Discussion

Model organisms’ power to elucidate general biological processes arises from the shared evolutionary ancestry of all organisms. However, evolutionary history is a double-edged sword, with lineage-specific changes limiting the general applicability of the mechanisms elucidated in model organisms. Distinguishing shared features of organisms from differentiated ones is a challenge, as is determining the ancestral (and thus perhaps more general) state. The study of the spliceosomal splicing is a case in point. On the one hand, remarkable progress has been made in understanding spliceosomal mechanisms in the yeast *S. cerevisiae*, much of which knowledge appears to apply quite well across a diversity of eukaryotes (e.g., (25)). On the other hand, comparative genomics of eukaryotes has revealed striking differences in the recognition of signals across species (e.g., (9,23,67). Given the centrality of *S. cerevisiae* for our understanding of splicing, observed qualitative general differences in splicing recognition signals between *S. cerevisiae* and most other eukaryotes are of particular importance. However, in the absence of a general framework for understanding these differences, it is challenging and arduous to identify and interpret the relevant differences in the splicing machinery and splicing signals.

15 years ago, Manuel Irimia spearheaded the first project to our knowledge offering a general approach to these differences. Studying splice signals across the diversity of eukaryotic lineages then available, we found evidence for evolutionary convergence of splicing signals that was unexpected, striking and strikingly predictable (9). Whereas in all species with many introns (humans, *S. pombe*, etc.) core splicing motifs were heterogeneous, consistent with intron relying on dispersed exonic and intronic motifs, in all species with few introns (including *S. cerevisiae*), core splicing motifs were homogeneous, consistent with intron recognition relying heavily on these short core motifs. Placement of these differences in phylogenetic context indicated that these differences are due to recurrent independent transformation of those lineages exhibiting few introns and heavy reliance on core splicing motifs, with order of magnitude intron number reduction and changes in intron recognition occurring independently in each of these lineages. The accumulation of genomic data has only underscored this picture, with lineages with few introns and concentrated splicing motifs emerging from within a background of more intron-rich species with heterogeneous splicing motifs (3,24,13–14,29,59,61).

These findings opened two different opportunities to address two very different larger questions. First, why do splicing mechanisms change in these transformed lineages? That is, why are changes in intron recognition mechanisms so closely associated with the genome-wide number or density of substrates? – a question with general implications for our understanding of biological signaling and the evolutionary forces acting on genomes. Second, how do splicing mechanisms change in these lineages? That is, what are the changes in splicing machinery that lead to the mechanistic changes in recognition, and how can knowledge of these changes improve our understanding of the differences and similarities in splicing mechanisms between different species?

To date, study of the evolutionary dynamics or splicing mechanisms in these transformed lineages has been limited. Analysis of the peculiar case of green algae from the group Mammieles, which harbor two different types of chromosomes with very different splicing signals, did not yield evidence for separate spliceosomal machineries, suggesting against our initial hypothesis of divergent selective pressures acting on splicing machineries with different numbers of substrates (since apparently the same machinery is responsible for two sets of introns using very different splicing signals; Irimia and Roy 2008). Study of the spliceosomal machinery in lineages with transformed splicing recognition have revealed a variety of peculiarities of the different lineages, including transformation of peripheral parts of snRNAs and loss and gain of spliceosomal components, including entire snRNPs (14,28–29,67). Here we report two studies of the changes in splicing machinery and evolutionary dynamics underlying changes in recognition machinery in one transformed lineage, *Entamoeba*.

### Complementary changes in snRNAs and splicing motifs

Complementary changes in interacting biomolecules is a recurrent theme in molecular and evolutionary biology. Interestingly, previous studies of species with atypical donor splice sites (in particular with pyrimidines at the 4^th^ position, e.g. ‘4Y’ changes) have mostly revealed typical U1 snRNAs that are thus not complementary to the most common donor splice site in the genome. For instance, 93% and 100% of introns in *S. cerevisiae* and in *Giardia lamblia* respectively exhibit a 4Y, and yet U1 snRNAs in both species retain the standard 3’-ACUUAC-5’ basepairing site, which is complementary to the standard GUAAGU (3,27–31). To our knowledge there is only one previous report of complementary changes of splice sites and U1 snRNAs, in the slime mold *Physarum polycephalum*, in which the most common donor splice sites is GUAUGU and in which reported U1 snRNAs show a complementary change (3’-ACAUAC-5’). However, further scrutiny of U1 snRNA candidates in this species in fact shows both standard and modified variants (these results will be presented in full elsewhere). Thus the case of *P. polycephalum* seems to be more complex.

In contrast, the current results show two independent cases of compensatory changes between a core spliceosomal snRNA and its target involving a total of three total changes in two related lineages (two in *Entamoeba* and one in *Mastigamoeba*). These results demonstrate that lack of complementary changes in other lineages does not reflect an impossibility of evolutionary change in core snRNA functions. Interestingly, previous results have suggested but were unable to conclusively demonstrate the importance for splicing of noncanonical snRNAs and complementary noncanonical splicing motifs in vertebrates (68,69).

### Cause and effect between selection, intron loss and intron transformation

Two hypotheses have been put forward to explain the association of homogeneous splice boundaries and intron paucity across species. First, the evolution of strict splicing requirements might lead to selection for loss of non-consensus introns (32). This hypothesis is of general interest for the general questions of crossing fitness valleys in evolution, since it posits that strict consensus requirements for splicing could evolve in the context of large numbers of nonconsensus introns, despite the fact that this renders introns with nonconsensus boundaries sufficiently costly they are then driven from the genome by selection. The second hypothesis holds that intron loss predates the evolution of strict consensus requirements for splicing, and that intron paucity allows for the evolution of strict consensus requirements, in ways that are not well understood (9,23).

The ongoing loss of introns from the genomes of *Entamoeba* species during relatively short evolutionary times allowed me to test whether nonconsensus introns are actually more likely to be lost in a lineage with homogenous splicing signals. We found no evidence that nonconsensus introns are more likely to be lost. Regardless of whether strict splicing requirements or intron loss occurs first evolutionarily, this result is somewhat surprising on its face. That most introns have consensus splice sites indicates that consensus splice sites are strongly favored by selection (since otherwise mutation would introduce heterogeneity), at least where they are observed. If consensus boundaries are generally preferred, this implies that introns with nonconsensus boundaries are more costly than are introns with consensus boundaries; thus random mutations deleting a nonconsensus intron should be more likely to succeed than random mutations deleting a consensus intron.

How can this paradox be resolved? We see two main possibilities. First, in *Entamoeba* all introns may be so costly that the difference in cost between consensus and nonconsensus introns does not substantially change the rate of evolutionary change. For instance, if nonconsensus boundaries only increase the cost of an intron by 10%, the probability of fixation of a mutation deleting a nonconsensus intron will only be roughly 10% higher than one deleting a consensus intron; evolutionary rates of loss will thus only differ by 10%, which would not produce significant results in a study such as this one. A second possibility is that the cost of non-consensus boundaries differs across sites. This could be the case if inefficient splicing was less costly in some genes (for instance, those with low expression), or if a substantial fraction of observed non-consensus boundaries played roles in gene expression regulation through splicing regulation (as seen in *S. cerevisiae* (70–71)). Whatever the explanation might be, the finding that nonconsensus introns are not preferentially lost from a lineage with homogeneous splice sites is in direct opposition to the predictions of the model of selective intron loss (32). However, given the small number of intron losses observed in ingroups in this genus, this result must be regarded as preliminary. Hopefully ongoing study will reveal species offering larger datasets of intron loss for analysis.

### Mechanisms of intron gain

Unexpectedly, comparison of intron positions across a broader diversity of amoebozoans revealed that a substantial fraction of the spliceosomal introns in *E. invadens* have been gained within the history of the genus. Such cases of recent large-scale gain of introns remain rare in the literature, and are of substantial interest. It is of note that substantial intron gain has occurred within the species of *Entamoeba*, since this genus exhibits two characteristics argued to be associated with efficient selection against introns (and thus low rates of intron creation). First, Lynch and Richardson (32) argued that species with strict requirements for core motifs (as in *Entamoeba*) are likely to shed and not accumulate introns, since these constraints impose increased mutational load on intron-containing alleles. Second, a variety of authors, arguing from different perspectives, have posited that the strong differences in intron numbers across species are driven by differences in the strength or efficiency of selection against introns (e.g., (72–73)). If this is so, then the same pressures that have driven intron numbers to low levels in organisms such as *Entamoeba* should also efficiently prevent the gain of new introns. Instead, intron gains in *E. invadens* appear to be abundant, accounting for a substantial fraction of introns in the genome. This represents the second instance in which a rare episode of widespread intron gain has been observed in an ancestrally intron-poor lineage (the other being in *Micromonas pusilla* (36)). These results are not as expected by models that rely on differences in selective strength or efficiency across lineages in determining the striking differences across lineages in genes and genome structures; instead, these results suggest that the availability of spontaneously occurring mutations producing changes in gene and genome structures may be a more important determining factor in the differences in evolutionary trajectory of genome structures across lineages (39,74).

The characteristics of the recently gained introns reported here provide a puzzle. On the one hand, the sequences of the recent intron gains show striking adherence to the strict splicing motifs shared across *Entamoeba* species, with gained introns showing equal adherence to consensus 5’ splice sites and branchpoint sequences. This striking degree of homogeneity is suggestive of a mechanism in which new introns are created by insertion of sequences already bearing splice site motifs. However, both proposed models that predict the presence of extended splicing motifs in new introns – insertion of transposable elements bearing splice site motifs (34,36–37) and movement of introns within the genome (75) – predict extended sequence similarity between introns in the genome, which is not observed here. Thus, if the extended splicing motifs observed in these newly-gained introns date to the origins of the introns, other parts of the introns must have since undergone substantial change to obscure the introns’ origins. On the other hand, if these introns were by and large created from sequences that were previously not associated with introns and thus not expected to bear extended motifs (e.g., (36,76–77)), these introns must have rapidly acquired these extended motifs. This requires reconciling the fixation of the initial intron-containing alleles, which would have had to be efficiently spliced despite lacking the extended splicing signals, with strong selection for the acquisition of these signals, presumably based on requirements for splicing efficiency. In total, then, the sequences of newly gained introns in *E. invadens* suggest rapid and complex evolution of recently created introns (35). Genomic sequencing of closer relatives of *E. invadens* could help to illuminate this intriguing case.

### Concluding remarks

These results expand our understanding of the dynamics by which eukaryotic genomes are transformed, the diversity of intron recognition machinery across eukaryotes, and the mechanisms by which introns are transformed. They also point to opportunities to bridge the gaps between the large literature on the mechanisms of splicing and the smaller but substantial literature on the comparative genomics of intron-exon structures across eukaryotes. A particularly profitable avenue of research may be comparative analyses of spliceosomal machinery in order to identify convergent changes taking place in the various transformed lineages, as these may pinpoint mechanistically important differences of the *S. cerevisiae* spliceosome from that of many other important eukaryotic species. Conceivably, such differences could provide targets for clinical interventions targeting the mechanistic differences in intron recognition between most eukaryotes and several parasites which exhibit transformed intron recognition – for instance *Cryptosporidium, Candida, Giardia, Trichomonas* and microsporidians.

## References

1. Nixon J.E., Wang A., Morrison H.G., McArthur A.G., Sogin M.L., Loftus B.J., Samuelson J. (2002) A spliceosomal intron in Giardia lamblia. Proc Natl Acad Sci U S A. 99, 3701–5.

2. Vanácová S., Yan W., Carlton J.M., Johnson P.J. (2005) Spliceosomal introns in the deep-branching eukaryote Trichomonas vaginalis. Proc Natl Acad Sci U S A. 102, 4430–5.

3. Roy S.W., Irimia M. (2014) Diversity and evolution of spliceosomal systems. Methods Mol Biol. 1126, 13–33.

4. Rogozin I.B., Carmel L., Csuros M., Koonin E.V. (2012) Origin and evolution of spliceosomal introns. Biol Direct. 7, 11.

5. Irimia M., Roy S.W. (2014) Origin of spliceosomal introns and alternative splicing. Cold Spring Harb Perspect Biol. , .

6. Lander E.S., Linton L.M., Birren B., Nusbaum C., Zody M.C., Baldwin J., Devon K., Dewar K., Doyle M., FitzHugh W., Funke R., Gage D., Harris K., Heaford A., Howland J., Kann L., Lehoczky J., LeVine R., McEwan P., McKernan K., Meldrim J., Mesirov J.P., Miranda C., Morris W., Naylor J., Raymond C., Rosetti M., Santos R., Sheridan A., Sougnez C., Stange-Thomann Y., Stojanovic N., Subramanian A., Wyman D., Rogers J., Sulston J., Ainscough R., Beck S., Bentley D., Burton J., Clee C., Carter N., Coulson A., Deadman R., Deloukas P., Dunham A., Dunham I., Durbin R., French L., Grafham D., Gregory S., Hubbard T., Humphray S., Hunt A., Jones M., Lloyd C., McMurray A., Matthews L., Mercer S., Milne S., Mullikin J.C., Mungall A., Plumb R., Ross M., Shownkeen R., Sims S., Waterston R.H., Wilson R.K., Hillier L.W., McPherson J.D., Marra M.A., Mardis E.R., Fulton L.A., Chinwalla A.T., Pepin K.H., Gish W.R., Chissoe S.L., Wendl M.C., Delehaunty K.D., Miner T.L., Delehaunty A., Kramer J.B., Cook L.L., Fulton R.S., Johnson D.L., Minx P.J., Clifton S.W., Hawkins T., Branscomb E., Predki P., Richardson P., Wenning S., Slezak T., Doggett N., Cheng J.F., Olsen A., Lucas S., Elkin C., Uberbacher E., Frazier M., Gibbs R.A., Muzny D.M., Scherer S.E., Bouck J.B., Sodergren E.J., Worley K.C., Rives C.M., Gorrell J.H., Metzker M.L., Naylor S.L., Kucherlapati R.S., Nelson D.L., Weinstock G.M., Sakaki Y., Fujiyama A., Hattori M., Yada T., Toyoda A., Itoh T., Kawagoe C., Watanabe H., Totoki Y., Taylor T., Weissenbach J., Heilig R., Saurin W., Artiguenave F., Brottier P., Bruls T., Pelletier E., Robert C., Wincker P., Smith D.R., Doucette-Stamm L., Rubenfield M., Weinstock K., Lee H.M., Dubois J., Rosenthal A., Platzer M., Nyakatura G., Taudien S., Rump A., Yang H., Yu J., Wang J., Huang G., Gu J., Hood L., Rowen L., Madan A., Qin S., Davis R.W., Federspiel N.A., Abola A.P., Proctor M.J., Myers R.M., Schmutz J., Dickson M., Grimwood J., Cox D.R., Olson M.V., Kaul R., Raymond C., Shimizu N., Kawasaki K., Minoshima S., Evans G.A., Athanasiou M., Schultz R., Roe B.A., Chen F., Pan H., Ramser J., Lehrach H., Reinhardt R., McCombie W.R., de la Bastide M., Dedhia N., Blöcker H., Hornischer K., Nordsiek G., Agarwala R., Aravind L., Bailey J.A., Bateman A., Batzoglou S., Birney E., Bork P., Brown D.G., Burge C.B., Cerutti L., Chen H.C., Church D., Clamp M., Copley R.R., Doerks T., Eddy S.R., Eichler E.E., Furey T.S., Galagan J., Gilbert J.G., Harmon C., Hayashizaki Y., Haussler D., Hermjakob H., Hokamp K., Jang W., Johnson L.S., Jones T.A., Kasif S., Kaspryzk A., Kennedy S., Kent W.J., Kitts P., Koonin E.V., Korf I., Kulp D., Lancet D., Lowe T.M., McLysaght A., Mikkelsen T., Moran J.V., Mulder N., Pollara V.J., Ponting C.P., Schuler G., Schultz J., Slater G., Smit A.F., Stupka E., Szustakowki J., Thierry-Mieg D., Thierry-Mieg J., Wagner L., Wallis J., Wheeler R., Williams A., Wolf Y.I., Wolfe K.H., Yang S.P., Yeh R.F., Collins F., Guyer M.S., Peterson J., Felsenfeld A., Wetterstrand K.A., Patrinos A., Morgan M.J., de Jong P., Catanese J.J., Osoegawa K., Shizuya H., Choi S., Chen Y.J., Szustakowki J; International Human Genome Sequencing Consortium. (2001) Initial sequencing and analysis of the human genome. Nature. 409, 860–921.

7. Mair G., Shi H., Li H., Djikeng A., Aviles H.O., Bishop J.R., Falcone F.H., Gavrilescu C., Montgomery J.L., Santori M.I., Stern L.S., Wang Z., Ullu E., Tschudi C. (2000) A new twist in trypanosome RNA metabolism: cis-splicing of pre-mRNA. RNA. 6, 163–9.

8. Slabodnick M.M., Ruby J.G., Reiff S.B., Swart E.C., Gosai S., Prabakaran S., Witkowska E., Larue G.E., Fisher S., Freeman RM J.r, Gunawardena J., Chu W., Stover N.A., Gregory B.D., Nowacki M., Derisi J., Roy S.W., Marshall W.F., Sood P. (2017) The Macronuclear Genome of Stentor coeruleus Reveals Tiny Introns in a Giant Cell. Curr Biol. 27, 569–575.

9. Irimia M., Penny D., Roy S.W. (2007) Coevolution of genomic intron number and splice sites. Trends Genet. 23, 321–5.

10. Warnecke T., Parmley J.L., Hurst L.D. (2008) Finding exonic islands in a sea of non-coding sequence: splicing related constraints on protein composition and evolution are common in intron-rich genomes. Genome Biol. 9, R29.

11. Wu X., Tronholm A., Cáceres E.F., Tovar-Corona J.M., Chen L., Urrutia A.O., Hurst L.D. (2013) Evidence for deep phylogenetic conservation of exonic splice-related constraints: splice-related skews at exonic ends in the brown alga Ectocarpus are common and resemble those seen in humans. Genome Biol Evol. 5, 1731–45.

12. Collins L., Penny D. (2005) Complex spliceosomal organization ancestral to extant eukaryotes. Mol Biol Evol. 22, 1053–66.

13. Hudson A.J., Stark M.R., Fast N.M., Russell A.G., Rader SD. Splicing diversity revealed by reduced spliceosomes in C. (2015) merolae and other organisms. RNA Biol. 12, 1–8.

14. Stark M.R., Dunn E.A., Dunn W.S., Grisdale C.J., Daniele A.R., Halstead M.R., Fast N.M., Rader S.D. (2015) Dramatically reduced spliceosome in Cyanidioschyzon merolae. Proc Natl Acad Sci U S A. 112, E1191–200.

15. Jurica, M. S., & Moore, M. J. (2003). Pre-mRNA splicing: awash in a sea of proteins. Molecular cell, 12(1), 5–14.

16. Rogozin I.B., Wolf Y.I., Sorokin A.V., Mirkin B.G., Koonin E.V. (2003) Remarkable interkingdom conservation of intron positions and massive, lineage-specific intron loss and gain in eukaryotic evolution. Curr Biol. 13, 1512–7.

17. Roy S.W., Gilbert W. (2005) Complex early genes. Proc Natl Acad Sci U S A. 102, 1986–91.

18. Roy S.W., Gilbert W. (2005) Rates of intron loss and gain: implications for early eukaryotic evolution. Proc Natl Acad Sci U S A. 102, 5773–8.

19. Carmel L., Wolf Y.I., Rogozin I.B., Koonin E.V. (2007) Three distinct modes of intron dynamics in the evolution of eukaryotes. Genome Res. 17, 1034–44.

20. Csuros M. (2005) Likely scenarios of intron evolution. RECOMB Workshop on Comparative Genomics. Springer Berlin Heidelberg, 2005.

21. Csuros M., Rogozin I.B., Koonin E.V. (2011) A detailed history of intron-rich eukaryotic ancestors inferred from a global survey of 100 complete genomes. PLoS Comput Biol. 7, e1002150.

22. Russell A.G., Charette J.M., Spencer D.F., Gray M.W. (2006) An early evolutionary origin for the minor spliceosome. Nature. 443, 863–6.

23. Irimia M., Roy S.W. (2008) Evolutionary convergence on highly-conserved 3’ intron structures in intron-poor eukaryotes and insights into the ancestral eukaryotic genome. PLoS Genet. 4, e1000148.

24. Xu F., Jerlström-Hultqvist J., Einarsson E., Astvaldsson A., Svärd S.G., Andersson J.O. (2014) The genome of Spironucleus salmonicida highlights a fish pathogen adapted to fluctuating environments. PLoS Genet. 10, e1004053.

25. Fabrizio P., Dannenberg J., Dube P., Kastner B., Stark H., Urlaub H., Lührmann R. (2009) The evolutionarily conserved core design of the catalytic activation step of the yeast spliceosome. Mol Cell. 36, 593–608.

26. Hudson, A. J., McWatters, D. C., Bowser, B. A., Moore, A. N., Larue, G. E., Roy, S. W., & Russell, A. G. (2019). Patterns of conservation of spliceosomal intron structures and spliceosome divergence in representatives of the diplomonad and parabasalid lineages. BMC Evolutionary Biology, 19(1).

27. Dávila López M., Rosenblad M.A., Samuelsson T. (2008) Computational screen for spliceosomal RNA genes aids in defining the phylogenetic distribution of major and minor spliceosomal components. Nucleic Acids Res. 36, 3001–10.

28. Kretzner L., Rymond B.C., Rosbash M. S. (1987) cerevisiae U1 RNA is large and has limited primary sequence homology to metazoan U1 snRNA. Cell. 50, 593–602.

29. Hudson A.J., Moore A.N., Elniski D., Joseph J., Yee J., Russell A.G. (2012) Evolutionarily divergent spliceosomal snRNAs and a conserved non-coding RNA processing motif in Giardia lamblia. Nucleic Acids Res. 40, 10995–1008.

30. Siliciano P.G., Jones M.H., Guthrie C. (1987) Saccharomyces cerevisiae has a U1-like small nuclear RNA with unexpected properties. Science. 237, 1484–7.

31. Simoes-Barbosa A., Meloni D., Wohlschlegel J.A., Konarska M.M., Johnson P.J. (2008) Spliceosomal snRNAs in the unicellular eukaryote Trichomonas vaginalis are structurally conserved but lack a 5’-cap structure. RNA. 14, 1617–31.

32. Lynch M., Richardson A.O. (2002) The evolution of spliceosomal introns. Curr Opin Genet Dev. 12, 701–10.

33. Collemare J., Beenen H.G., Crous P.W., de Wit P.J., van der Burgt A. (2015) Novel Introner-Like Elements in fungi Are Involved in Parallel Gains of Spliceosomal Introns. PLoS One. 10, e0129302.

34. Huff J.T., Zilberman D., Roy S.W. (2016) Mechanism for DNA transposons to generate introns on genomic scales. Nature. 538, 533–536.

35. Li W., Tucker A.E., Sung W., Thomas W.K., Lynch M. (2009) Extensive, recent intron gains in Daphnia populations. Science. 326, 1260–2.

36. Worden A.Z., Lee J.H., Mock T., Rouzé P., Simmons M.P., Aerts A.L., Allen A.E., Cuvelier M.L., Derelle E., Everett M.V., Foulon E., Grimwood J., Gundlach H., Henrissat B., Napoli C., McDonald S.M., Parker M.S., Rombauts S., Salamov A., Von Dassow P., Badger J.H., Coutinho P.M., Demir E., Dubchak I., Gentemann C., Eikrem W., Gready J.E., John U., Lanier W., Lindquist E.A., Lucas S., Mayer K.F., Moreau H., Not F., Otillar R., Panaud O., Pangilinan J., Paulsen I., Piegu B., Poliakov A., Robbens S., Schmutz J., Toulza E., Wyss T., Zelensky A., Zhou K., Armbrust E.V., Bhattacharya D., Goodenough U.W., Van de Peer Y., Grigoriev I.V. (2009) Green evolution and dynamic adaptations revealed by genomes of the marine picoeukaryotes Micromonas. Science. 324, 268–72.

37. van der Burgt A., Severing E., de Wit P.J., Collemare J. (2012) Birth of new spliceosomal introns in fungi by multiplication of introner-like elements. Curr Biol. 22, 1260–5.

38. Roy S.W., Fedorov A., Gilbert W. (2003) Large-scale comparison of intron positions in mammalian genes shows intron loss but no gain. Proc Natl Acad Sci U S A. 100, 7158–62.

39. Roy S.W., Hartl D.L. (2006) Very little intron loss/gain in Plasmodium: intron loss/gain mutation rates and intron number. Genome Res. 16, 750–6.

40. Ma M.Y., Che X.R., Porceddu A., Niu D.K. (2015) Evaluation of the mechanisms of intron loss and gain in the social amoebae Dictyostelium. BMC Evol Biol. 15, 286.

41. Ma M.Y., Zhu T., Li X.N., Lan X.R., Liu H.Y., Yang Y.F., Niu D.K. (2015) Imprecise intron losses are less frequent than precise intron losses but are not rare in plants. Biol Direct. 10, 24.

42. Stajich J.E., Dietrich F.S. (2006) Evidence of mRNA-mediated intron loss in the human-pathogenic fungus Cryptococcus neoformans. Eukaryot Cell. 5, 789–93.

43. Yang Y.F., Zhu T., Niu D.K. (2013) Association of intron loss with high mutation rate in Arabidopsis: implications for genome size evolution. Genome Biol Evol. 5, 723–33.

44. Zhu T., Niu D.K. (2013) Mechanisms of intron loss and gain in the fission yeast Schizosaccharomyces. PLoS One. 8, e61683.

45. Roy S.W., Penny D. (2007) A very high fraction of unique intron positions in the intron-rich diatom Thalassiosira pseudonana indicates widespread intron gain. Mol Biol Evol. 24, 1447–57.

46. Farhat, S., Le, P., Kayal, E., Noel, B., Bigeard, E., Corre, E., Maumus, F., Florent, I., Alberti, A., Aury, J.M. and Barbeyron, T. (2021). Rapid protein evolution, organellar reductions, and invasive intronic elements in the marine aerobic parasite dinoflagellate Amoebophrya spp. BMC biology, 19(1), pp.1–21.

47. Simmons, M.P., Bachy, C., Sudek, S., Van Baren, M.J., Sudek, L., Ares Jr, M. and Worden, A.Z. (2015) Intron invasions trace algal speciation and reveal nearly identical Arctic and Antarctic Micromonas populations. Molecular biology and evolution, 32(9), pp.2219–2235.

48. Henriet, S., Sanmartí, B.C., Sumic, S. and Chourrout, D., 2019. Evolution of the U2 spliceosome for processing numerous and highly diverse non-canonical introns in the chordate Fritillaria borealis. Current Biology, 29(19), pp.3193–3199.

49. Davis C.A., Brown M.P., Singh U. (2007) Functional characterization of spliceosomal introns and identification of U2, U4, and U5 snRNAs in the deep-branching eukaryote Entamoeba histolytica. Eukaryot Cell. 6, 940–8.

50. Roy S.W., Irimia M., Penny D. (2006) Very little intron gain in Entamoeba histolytica genes laterally transferred from prokaryotes. Mol Biol Evol. 23, 1824–7.

51. Roy, S.W., Penny D. (2007) Intron length distributions and gene prediction. Nucleic Acids Res 14:4737–42.

52. Crooks G.E., Hon G., Chandonia J.M., Brenner S.E. (2004) WebLogo: a sequence logo generator. Genome Res. 14, 1188–90.

53. Griffiths-Jones S., Bateman A., Marshall M., Khanna A., Eddy S.R. (2003) Rfam: an RNA family database. Nucleic Acids Res. 31, 439–41.

54. Altschul, S.F., Madden, T.L., Schäffer, A.A., Zhang, J., Zhang, Z., Miller, W. & Lipman, D.J. (1997) Gapped BLAST and PSI-BLAST: a new generation of protein database search programs. Nucleic Acids Res. 25, 3389–3402.

55. Johnson L. S., Eddy S. R., & Portugaly E. (2010). Hidden Markov model speed heuristic and iterative HMM search procedure. BMC Bioinformatics, 11(1), 431.

56. Tatusov R. L., Koonin E. V, & Lipman D. J. (1997). A Genomic Perspective on Protein Families. Science, 278(5338), 631 LP – 637.

57. Bork, P., Dandekar, T., Diaz-Lazcoz, Y., Eisenhaber, F., Huynen, M., & Yuan, Y. (1998). Predicting function: from genes to genomes and back. Edited by P. E. Wright. Journal of Molecular Biology, 283(4), 707–725.

58. Russell A.G., Shutt T.E., Watkins R.F., Gray M.W. (2005) An ancient spliceosomal intron in the ribosomal protein L7a gene (Rpl7a) of Giardia lamblia. BMC Evol Biol. 5, 45.

59. Lee R.C., Gill E.E., Roy S.W., Fast N.M. (2010) Constrained intron structures in a microsporidian. Mol Biol Evol. 27, 1979–82.

60. Du H., Rosbash M. (2002) The U1 snRNP protein U1C recognizes the 5’ splice site in the absence of base pairing. Nature. 419, 86–90.

61. Derelle E., Ferraz C., Rombauts S., Rouzé P., Worden A.Z., Robbens S., Partensky F., Degroeve S., Echeynié S., Cooke R., Saeys Y., Wuyts J., Jabbari K., Bowler C., Panaud O., Piégu B., Ball S.G., Ral J.P., Bouget F.Y., Piganeau G., De Baets B., Picard A., Delseny M., Demaille J., Van de Peer Y., Moreau H. (2006) Genome analysis of the smallest free-living eukaryote Ostreococcus tauri unveils many unique features. Proc Natl Acad Sci U S A. 103, 11647–52.

62. Farlow A., Meduri E., Dolezal M., Hua L., Schlötterer C. (2010) Nonsense-mediated decay enables intron gain in Drosophila. PLoS Genet. 6, e1000819.

63. Farlow A., Meduri E., Schlötterer C. (2011) DNA double-strand break repair and the evolution of intron density. Trends Genet. 27, 1–6.

64. Sullivan, J. C., Reitzel, A. M., & Finnerty, J. R. (2006). A High Percentage of Introns in Human Genes Were Present Early in Animal Evolution Evidence from the Basal Metazoan Nematostella vectensis. Genome Informatics, 17(1), 219–229.

65. Roy, S.W. (2016) Probing the mechanisms of intron creation in a fast-evolving mite. biorxiv doi.org/10.11101/051292.

66. Sverdlov, A. V., Rogozin, I. B., Babenko, V. N., & Koonin, E. V. (2003). Evidence of splice signal migration from exon to intron during intron evolution. Current Biology, 13(24), 2170–2174.

67. Schwartz S.H., Silva J., Burstein D., Pupko T., Eyras E., Ast G. (2008) Large-scale comparative analysis of splicing signals and their corresponding splicing factors in eukaryotes. Genome Res. 18, 88–103.

68. O’Reilly D., Dienstbier M., Cowley S.A., Vazquez P., Drozdz M., Taylor S., James W.S., Murphy S. (2013) Differentially expressed, variant U1 snRNAs regulate gene expression in human cells. Genome Res. 23, 281–91.

69. Kyriakopoulou C., Larsson P., Liu L., Schuster J., Söderbom F., Kirsebom L.A., Virtanen A. (2006) U1-like snRNAs lacking complementarity to canonical 5’ splice sites. RNA. 12, 1603–11.

70. Davis C.A., Grate L., Spingola M., Ares M J.r (2000) Test of intron predictions reveals novel splice sites, alternatively spliced mRNAs and new introns in meiotically regulated genes of yeast. Nucleic Acids Res. 28, 1700–6.

71. Eng, F. J., & Warner, J. R. (1991). Structural basis for the regulation of splicing of a yeast messenger RNA. Cell, 65(5), 797–804.

72. Doolittle, W. F. (1978). Genes in pieces: were they ever together?. Nature, 272(5654), 581–582.

73. Lynch, M., & Richardson, A. O. (2002). The evolution of spliceosomal introns. Current opinion in genetics & development, 12(6), 701–710.

74. Collemare J., van der Burgt A., de Wit P.J. (2013) At the origin of spliceosomal introns: Is multiplication of introner-like elements the main mechanism of intron gain in fungi. Commun Integr Biol. 6, e23147.

75. Denoeud F., Henriet S., Mungpakdee S., Aury J.M., Da Silva C., Brinkmann H., Mikhaleva J., Olsen L.C., Jubin C., Cañestro C., Bouquet J.M., Danks G., Poulain J., Campsteijn C., Adamski M., Cross I., Yadetie F., Muffato M., Louis A., Butcher S., Tsagkogeorga G., Konrad A., Singh S., Jensen M.F., Huynh Cong E., Eikeseth-Otteraa H., Noel B., Anthouard V., Porcel B.M., Kachouri-Lafond R., Nishino A., Ugolini M., Chourrout P., Nishida H., Aasland R., Huzurbazar S., Westhof E., Delsuc F., Lehrach H., Reinhardt R., Weissenbach J., Roy S.W., Artiguenave F., Postlethwait J.H., Manak J.R., Thompson E.M., Jaillon O., Du Pasquier L., Boudinot P., Liberles D.A., Volff J.N., Philippe H., Lenhard B., Roest Crollius H., Wincker P., Chourrout D. (2010) Plasticity of animal genome architecture unmasked by rapid evolution of a pelagic tunicate. Science. 330, 1381–5.

76. Curtis B.A., Archibald J.M. (2010) A spliceosomal intron of mitochondrial DNA origin. Curr Biol. 20, R919–20.

77. Hellsten U., Aspden J.L., Rio D.C., Rokhsar D.S. (2011) A segmental genomic duplication generates a functional intron. Nat Commun. 2, 454.

78. Kang S., Tice F.T., Spiegel F.W., Silberman J.D., Pánek T., Čepička I., Kostka M., Kosakyan A., Alcântara DMC., Roger A.J., Shadwick L.L., Smirnov A., Kudryavtsev A., Lahr D.J.G., Brown M.W., Between a Pod and a Hard Test: The Deep Evolution of Amoebae, Molecular Biology and Evolution, Volume 34, Issue 9, September 2017, 2258–2270.

